# Creativity Potential Networks: Brain Markers for Novelty and Feasibility of Upcoming Divergent Thinking Solutions

**DOI:** 10.1101/2025.06.24.661027

**Authors:** Yiyuan Teresa Huang, Chien-Te Wu, Zenas C. Chao

## Abstract

The brain’s resting-state activity can serve as an indicator of cognitive flexibility and predict the likelihood of an upcoming Aha experience. This suggests that spontaneous neural dynamics reflect a person’s readiness for creative insight and underscore the potential of resting-state measures as biomarkers for anticipating creative breakthroughs. However, solutions accompanied by an Aha experience are not always truly creative, so it may be more valuable to identify biomarkers specifically linked to novelty and usefulness—two key dimensions of creative performance. To achieve this, we recruit 49 participants to complete the Alternative Uses Test, in which unconventional uses for everyday items are generated. We evaluate the responses for both novelty and feasibility using automated GPT-based methods and analyze resting-state EEG prior to the test. We find that creative performance is better predicted by interactions between different brain areas than by the activation of individual regions. Specifically, the degree centrality of theta-band functional connectivity in the right parietal and occipital areas correlates with novelty, while connectivity in the right middle and inferior frontal areas is associated with more feasible answers. These findings highlight distinct resting-state brain networks underlying the “creative potential” for novelty and feasibility, which could be leveraged to monitor and enhance brain flexibility.

**Significance statement:** Our study introduces the Creativity Potential Network (CPN), a resting-state brain network that can predict the novelty and feasibility of the upcoming solution in creative problem-solving. We show that the CPN is represented by communication between brain areas, and that the networks for novelty and feasibility are spatially distinct. This work provides a potential method to assess the potential to be creative without relying on behavioral measures and could be combined with neurofeedback to monitor and enhance brain flexibility.

## Introduction

Creativity, defined as the capacity to generate novel and useful solutions, has been supported by flexibility of the brain state (Beaty et al., 2015, 2018; Chen et al., 2025; Fink et al., 2009; Sun et al., 2019). That is, when the brain during or prior to solving problems is more flexible and connected widespread, more creative solutions could be obtained. For example, in the Remote Associates Test (RAT)— a convergent thinking task— participants are presented with three words (e.g., apple, house, family) and required to find a word that can form a compound or phrase with each of the given words (e.g., “tree”). Brain activity preceding the question, particularly alpha power in EEG and BOLD activation in the middle and superior temporal and cingulate regions, predicts the occurrence of the “Aha” moment— a sudden insight that leads to a creative solution (Kounios et al., 2006). However, subjective Aha experience can also be accompanied with incorrect solutions, known as false insights (Danek & Wiley, 2017), and may even bias the evaluation of solutions (Laukkonen et al., 2020). Therefore, an Aha experience alone may not definitively indicate true insight, and other phenomenological and behavioral variables might be needed (Danek & Wiley, 2017).

To examine whether brain activity prior to the task presentation contains information about upcoming creativity performance (referred to as “creativity potential”), we focus on more objective measures of creativity performance, particularly the novelty and feasibility of the solutions. These measures stem directly from the definition of creativity—the ability to generate original and useful ideas. To achieve this, we use the Alternative Uses Test (AUT), a divergent thinking task shown to predict real-world creativity (Kim, 2008; Said-Metwaly et al., 2024; Snyder et al., 2019), and which allows for the evaluation of both the novelty and feasibility of a solution. In the test, participants are asked to generate alternative uses for an everyday item (e.g., using an umbrella as a boat). To assess the novelty and feasibility of solutions in the AUT, we use a validated and automated GPT-based method that provides evaluations comparable to those of a cohort of human judges (Kern et al., 2024). Also, to provide a better spatial view of the creativity potential response, we record EEG signals and then interpolate the signals into the brain cortex using the standard MR images. We then measure brain activation in individual regions and inter-regional interactions during the resting period preceding each trial, and compare these between high- and low-novelty trials, and between high- and low-feasibility trials. We find significant differences in inter-regional interactions—but not in regional activations—between high and low performance, suggesting that the potential to generate creative solutions depends on communication between brain regions. We refer to these significant networks as the “Creativity Potential Networks” (CPNs). Interestingly, the CPNs associated with novelty and feasibility are distinct, indicating that different neural mechanisms underlie these two key elements of creative thinking.

## Results

### Alternative Uses Test (AUT)

To investigate how novelty and feasibility are influenced by the preparatory brain state, we recruited 49 participants (16 males and 33 females, 24.2 ± 3.6 years old) to perform a Japanese Alternative Uses Test (AUT), while their brain signals recorded using a 128-channel EEG system. The test consisted of 30 trials, each presenting an everyday item as the question (see Supplementary Table 1). Participants were instructed to press a button as soon as they thought of an unconventional use for the item (see Figure 1 and details in Methods), and then provide a self-reported “Aha” rating on a scale from 1 (weak) to 7 (strong). Reaction times were recorded, with an average 15.0 ± 9.2 seconds (mean ± standard deviation, n = 49 participants). The novelty and feasibility scores of each solution were evaluated using an automated GPT-based method (Kern et al., 2024).

**Figure 1.**
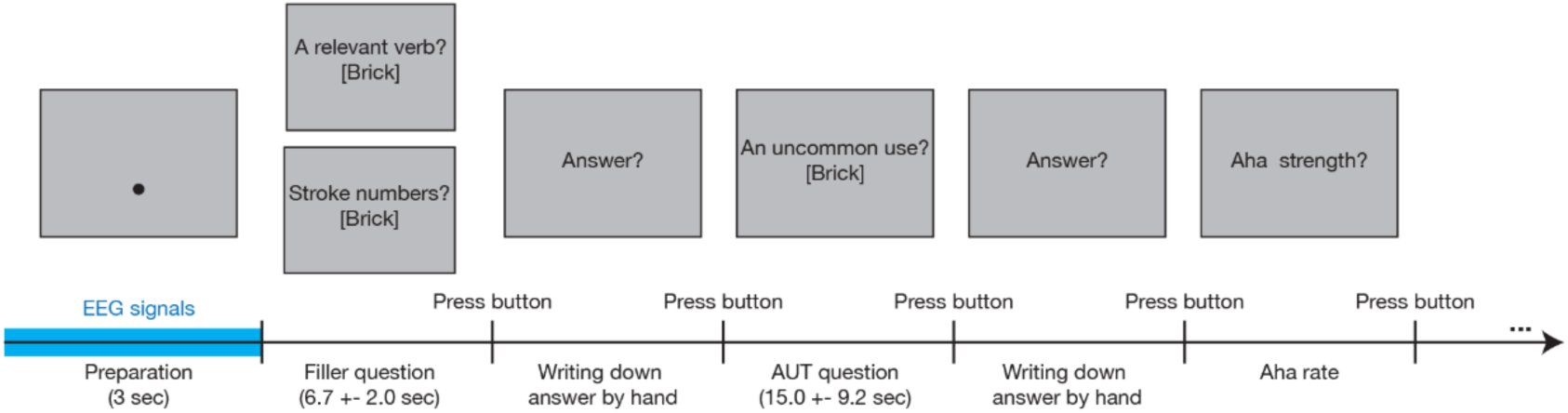
**Task design**

We found a significant negative correlation between the novelty and feasibility scores (*p* < 2e-16, beta = -0.745, see Supplementary Figure 1A) (n = 49 participants, linear mixed model, random effect = the participant, alpha = 0.05), suggesting more novel solutions tend to be less feasible, and vice versa. It is worth noting that AUT produces solutions which cannot be simply categorized into accurate or inaccurate ones; therefore, we cannot evaluate false “Aha”. However, we found “Aha” ratings were correlated with the novelty scores (*p* = 0.013, estimated coefficient β = 0.003) (Supplementary Figure 1B) but not with the feasible scores (*p* = 0.201, estimated coefficient β = -0.001) (Supplementary Figure 1C). This indicates a stronger feeling of insight is associated with more novel solutions but not necessarily with greater feasibility.

## Creativity potential represented in functional connectivity but not power

To extract preparatory responses related to the novelty and feasibility of upcoming solutions, we analyzed EEG signals during the 3 seconds resting-period preceding stimulus presentation (highlighted in blue in Figure 1). For each participant, we categorized the 30 trials into two conditions: *high novelty* and *low novelty*, based on the median novelty score for each participant (59.44 ± 14.62; mean ± standard deviation across participants). The same categorization was done for feasibility scores, dividing them into *high feasibility* and *low feasibility* (38.61 ± 20.71). Then, EEG signals were processed to reconstruct activities at 4050 source nodes (see details in Methods).

We estimated the power and coherence of the source signals across four frequency bands: theta (4-8 Hz), alpha (8-13 Hz), beta (13-30 Hz), and gamma (30-50 Hz). For the power analysis, we projected the power from the source nodes to the cortical surface (116 parcels), with the anatomical parcellation (see details in Methods). For the coherence analysis, we estimated degree centrality, which quantifies the number of functional connections between a specific source node and all other source nodes whose connectivity exceeds a predetermined threshold. In other words, degree centrality measures the extent to which a source is interconnected with other sources. Connections were retained or discarded based on whether their coherence value exceeded or fell below the threshold. To address the arbitrariness of threshold selection, we tested three values (0.05, 0.1, and 0.15) to ensure the robustness of subsequent analyses. After computing the degree centrality for the source nodes, the results were projected onto the cortical surface, with each of the 116 parcels assigned a single degree centrality value.

Then, we compared the power values between *high novelty* and *low novelty* trials and between *high feasibility* and *low feasibility* trials using the Monte Carlo method (1,000 randomizations, alpha = 0.05, two-tailed, false discovery rate). Similar comparisons were conducted for the degree centrality values. First, we found no significant differences in power across the four frequency bands when comparing *high novelty* and *low novelty* trials, as well as between *high feasibility* and *low feasibility* trials. In contrast, significant differences in degree centrality were observed in the theta frequency band for both novelty and feasibility (Figure 2), representing the Creativity Potential Networks (CPNs) for novelty and feasibility, respectively. With the threshold of 0.1, the CPN for novelty involved the left inferior frontal area, right superior and inferior parietal areas, right angular gyrus, right superior and middle occipital areas, and right cuneus (Figure 2A). On the other hand, the CPN for feasibility involved the left superior temporal area, right insula, left Rolandic operculum, left middle occipital area, and right middle and inferior frontal areas (Figure 2B). Notably, there were no overlapping brain areas between these two CPNs. A consistent CPN across different threshold selections was found only in the theta band (see results for thresholds of 0.05 and 0.15 in Supplementary Figures 2 and 3).

**Figure 2.**
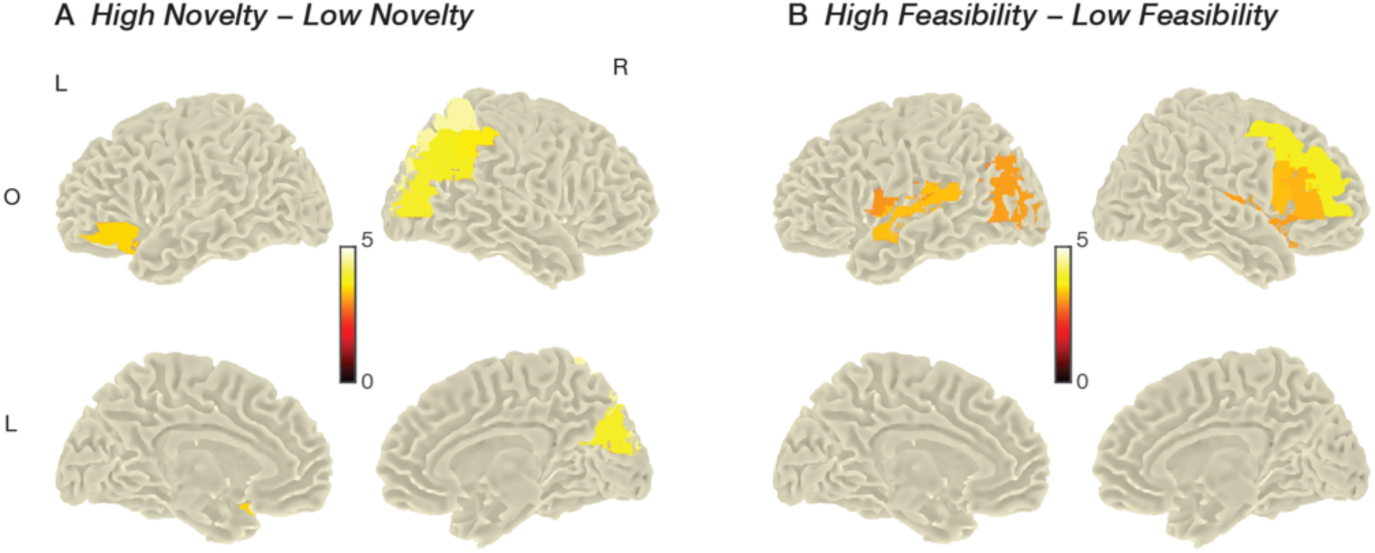
Functional connectivity density during preparation. The color bars represent statistical values.

## Discussion

We combined the commonly used AUT with GPT-based evaluation to examine creativity potential in the brain. Our findings demonstrated that creativity potential is reflected in theta-band connectivity between brain areas, and not in the singular brain activations. Importantly, distinct brain areas were identified as key hubs for novelty and feasibility. To our knowledge, this is the first study investigating how pre-task brain states predict the novelty and feasibility of creative performance.

### Creativity potential in distributed network communication

Our research reveals that the creativity potential is represented through functional connectivity in the brain, rather than being confined to specific regions. We suggest that the pattern of distributed brain activations is crucial for divergent thinking. During the AUT, achieving more creative performance often involves expanding the search and disregarding trivial solutions. This process likely engages distributed brain networks to enable a wider array of potential solutions. This is evidenced by findings that distributed brain regions, including the anterior cingulate, inferior frontal, temporal, and parietal areas, are activated during the search for solutions (Fink et al., 2009; Wu et al., 2015). In addition, our finding that stronger connectivity is associated with better creative performance aligns with existing literature showing a link between greater resting-state connectivity and improved performance on creativity-related questionnaires or tasks outside the MRI scanner. For example, increased connectivity between the inferior frontal gyrus and the default mode network (DMN) has been observed in individuals with high creativity, as assessed by a battery of divergent thinking tests (Beaty et al., 2014). The DMN is typically considered a task-negative network, becoming more active when no specific task is being performed, and it has been associated with mind wandering and spontaneous thought (Fox et al., 2015; Poerio et al., 2017). In contrast, the executive control network (ECN) is a task-positive network, commonly engaged in response control and task switching (Hwang et al., 2010; Menon & D’Esposito, 2022). Notably, while stronger activity within the DMN has been associated with creativity, the coupling between the DMN and ECN is gaining increasing attention in the field (Beaty et al., 2015, 2018; Ellamil et al., 2012). Specifically, stronger coupling between these networks has been observed in individuals with higher creativity, and their synchronization is thought to reflect the complementary processes of exploration (DMN) and evaluation (ECN) (Beaty et al., 2018). Importantly, while prior studies focus on differences between creative and non-creative individuals, we examine differences between flexible and rigid states within each individual. This approach allows us to monitor creative potential over time and opens new avenues for intervention. It is worth noting that our power analysis showed no significant results, which contrasts with the findings of Kounios et al. (2016) based on the RAT. While Aha experiences in the RAT have been linked to long-distance search (Chao et al., 2025), the RAT, as a verbal puzzle, likely restricts the search range to the semantic domain and engages fewer dimensions than the domain-general AUT.

### Distinct brain area hubs for the novelty and feasibility

Our second key finding demonstrates that novelty and feasibility can be predicted by connectivity to distinct active brain hubs. Specifically, novelty is associated with a higher degree centrality primarily in the right superior and inferior parietal areas, and thus we infer the preparatory process of novelty may involve the DMN. This network has been reported to include the posterior cingulate cortex, precuneus, and inferior parietal regions (Broyd et al., 2009; Raichle et al., 2001). The inferior parietal area, as a key component of the DMN, was identified in a functional magnetic resonance imaging (fMRI) study where increased activation was observed when comparing new ideas to old ones in the AUT (Fink et al., 2010). Additionally, research using a word drawing test showed increased activation in the parietal cortex when drawing a new word compared to a control word (Saggar et al., 2015). On the other hand, we found that feasibility is associated with a higher degree centrality mostly in the right middle and inferior frontal areas, which are core components of the ECN (Thomas Yeo et al., 2011). The frontal areas within this network have also been identified in various creative processes (Gonen-Yaacovi et al., 2013). For instance, music creation involves activation in the dorsolateral prefrontal cortex as well as motor areas (Bengtsson et al., 2007). A creative solution should be both novel and feasible, as demonstrated by the higher degree centrality for novelty and feasibility associated with the DMN and the ECN, respectively. Although considerable research indicates these two networks operate in an opposite manner, e.g., DMN activates but the ECN deactivates during the rest, the coupling of the networks has growing attention, especially in the creativity field, where greater co-activations are associated with higher creativity (Beaty et al., 2015, 2018). Our study further provides interpretations of the crucial coupling because these two networks are responsible for different components of creativity.

### Differences in brain areas compared to previous studies

Literature reviews and studies have revealed a hemispheric specialization of creativity predominantly in the left hemisphere (Aziz-Zadeh et al., 2013; Saggar et al., 2015), while our findings show the higher connection to the right parietal areas, particularly for novelty. This discrepancy could be attributed to the different time periods we focused on in our analyses.

Previous studies often used task-related responses, whereas our approach focused on responses during resting states. This can be supported by a study, where Beaty and his colleagues (2015) identified the right dorsolateral prefrontal cortex, a component of the DMN, as a seed region, and found a stronger connection to the right inferior parietal lobe. Furthermore, while we reconstructed source signals based on a structural standard brain image to enhance EEG spatial resolution, incorporating data from acquisition modalities with higher spatial resolution is necessary to better connect our findings with existing literature and provide significant implications for brain stimulation aimed at enhancing novelty and feasibility. In addition, we adopted a data-driven approach and did not conduct seed-based functional connectivity analysis targeting specific brain areas. In this analysis, two commonly used metrics are degree centrality and betweenness centrality, the latter evaluated as a measure of a bridge node between two nodes based on the shortest path. We also calculated betweenness centrality, but the results were inconclusive. Specifically, the threshold parameter in network analysis is arbitrary, and varying threshold values can yield different brain results across different frequency bands (see Supplementary Figures 4 and 5). We attribute this variability to the short duration of resting state recording (3 seconds), and suggest that longer trial lengths will be needed to address this.

### Theta oscillation

We demonstrate that both novelty and feasibility can be predicted by theta-band coherence. This slow oscillation, particularly in the frontal region, has been recognized as playing a significant role in cognitive control (Cavanagh & Frank, 2014). For instance, frontal midline theta (FMθ) activity has been shown to increase following the presentation of novel or conflicting stimuli, suggesting that enhanced cognitive control is engaged to optimize performance in subsequent trials (Cavanagh et al., 2012). Furthermore, the theta oscillation has been suggested to facilitate collaboration among distinct brain areas, creating a task-related network (Fries, 2015). Rhythmic transcranial magnetic stimulation (TMS) at theta frequency has been shown to improve memory capacity and reaction speed (Riddle et al., 2020). Additionally, a neural synchrony model suggests that a random burst in the theta frequency enhances behavioral performance in the Stroop cognitive control task and modulates communication with posterior brain regions (Verguts, 2017). In creativity research, increased delta and theta coherence has been observed when participants created a short story incorporating preassigned words (Petsche, 1996). Also, by increasing the ratio of alpha and theta waves through auditory stimuli, an EEG-neurofeedback protocol has been found to improve creativity (Boynton, 2001; Gruzelier, 2009). These findings support our observation of the frontal theta as a central hub, indicating that stronger theta-band connections between brain areas enable flexible shifts toward task targets and enhance creative performance.

In summary, our work identifies brain markers for the potential to generate novel and feasible solutions in divergent thinking, paving the way for neurofeedback systems that enable users to train themselves to become more flexible in their thinking.

## Methods

### Participants

Forty-nine participants were recruited, following the protocols approved by the Research Ethics Committee at The University of Tokyo (No. 21-380). All the participants were native Japanese speakers and had no difficulty in understanding the experimental procedure due to apparent deficits in cognition, hearing and vision and no medical history of neurological or psychological diagnosis. Signed consent was received from the participants before the experiment.

### AUT and its evaluation

For the test, we presented 30 daily items (see all items in Supplementary Table 1) randomly for each participant. Each trial began with a 3-second preparation where the participant was only asked to fixate on a black dot on the monitor for controlling eye movements during EEG recording (position: 120 pixels below the center of the monitor). Then, a filler question was presented, asking either a verb that is associated with an item (e.g., “hold” an umbrella) or stroke numbers (e.g., 12 strokes for umbrella in kanji or katakana). The participant pressed a button and wrote down their answer on the tablet, and there was no time limitation. The average reaction time across the participants is 6.7 +-2.0 second. Those two types of filler questions were demonstrated with an equal number (i.e., 15 trials for asking verbs and 15 trials for asking stroke numbers) and in a random order.

After the filler question, the same item was presented. The participant was required to come up with one alternative use of the item as soon as possible and respond by pressing a button (no time limitation). The average reaction time is 15.0 ± 9.2 second, and the answer was written down on a tablet. Then, they were also asked to score the strength of their “Aha” feeling when they found out their answer for the unusual use with a 7-point Likert Scale from one (no “Aha”) to seven (strong “Aha”). Please that we designed filler questions for other purposes, but there are no significant differences between the two types of the filler question on “Aha”, novelty, and feasibility values (*p* = 0.377, 0.473, and 0.288 respectively). The correlations among three kinds of the score was estimated using linear mixed model with the participant as the random effect for controlling individual differences (R function: *lmer*).

The stimulation delivery was done using MATLAB-based PsychtoolBox (Kleiner et al., 2007; Pelli, 1997). We set the gray color for the background of the monitor (RGB: [200, 200, 200]). All the instructions and questions were delivered in Japanese font and black color.

### EEG recording and analysis

EEG raw data was recorded by the actiCHamp Plus amplifier and 128-channel actiCAP slim system (Brain Product, Inc.). The impedances were kept below 30 kΩ, and the data were collected with an online reference electrode Cz and a 1000-Hz sampling rate. We used EEGLAB (Swartz Center for Computational Neuroscience) (Delorme & Makeig, 2004) for data preprocessing for each participant. First, the data was filtered with a 0.05-Hz high-pass filter (function: pop_eegfiltnew.m). We applied independent component analysis (ICA) (function: pop_ica.m) and removed artificial components with the ADJUST toolbox (Mognon et al., 2011) (function: interface_ADJ.m). The processed data were then resampled to a 250-Hz rate and re-reference the average of all electrodes. Because of the focus on preparatory responses, we extracted epochs from -3 second to 0 relative to the onset of each trial.

### Source reconstruction analysis

We used MATLAB-based Fieldtrip (Oostenveld et al., 2010) to obtain two main measurements: power and functional connectivity across four frequency bands (theta: 4-8 Hz; alpha: 8-13 Hz; beta: 13-30 Hz; gamma: 30-50 Hz), and then transformed both from the channel-level to the source level. First, for each participant, we grouped the processed data into *high novelty* and *low novelty* based on the median of novelty values and into *high feasibility* and *low feasibility* based on the median of feasibility values. Then, we estimated the frequency representations of those five datasets, including the whole dataset (function: *ft_freqanalysis.mat*). Second, for the source analysis, we used a standard head model (also known as a volume conduction model), created by an averaged magnetic resonance image (MRI) data from Montreal Neurological Institute and Boundary Element Method (filename: *standard_bem.mat*). The electrode location was aligned to the head model, and leadfield (potential contributions from dipoles to electrodes) was generated with the head model and the electrode location, with a resolution of 1 center (function: *ft_prepare_leadfield.m*).

For the power at the source level and for each participant, we applied the dynamic imaging of coherent sources method first to the whole dataset to extract a common spatial filter. Then, source responses of the four separate datasets were estimated using this filter (function: *ft_sourceanalysis.m*). The source-level power in each of the four datasets was interpolated and parcellated based on the AAL atlas (with 116 brain regions) (functions: *ft_sourceinterpolate.m*, *ft_sourceparcellate.m*).

For the degree centrality of functional connectivity and for each participant, we applied the partial canonical coherence method, and the same procedure for the whole dataset and then the others were done. Then, we calculated the imaginary part of coherence of the source signals in the four datasets (function: *ft_connectivityanalysis.m*) and degrees with the thresholds: 0.05, 0.1, and 0.15, respectively (function: *ft_networkanalysis.m*). Finally, the source-level degree in each of the datasets was interpolated and parcellated based on the AAL atlas.

We estimated if the EEG response during preparation is significantly different from zero between more and less novel performances and between more and less feasible performances. For each frequency band in the power and degree and each of the comparisons between *high novelty* and *low novelty* and between *high feasibility* and *low feasibility*, significances across participants were tested using the Monte Carlo method with parameters set to 1000 randomizations, an alpha of 0.05, two tails, and false discovery rate for the alpha correction (function: *ft_sourcestatistics.m)*. The significances were visualized on four cortical surfaces of the MNI brain (left and right hemispheres, and outside and inside views), with parameters set to the nearest projection and no lighting (*ft_sourceplot.m*).

## Data availability

The raw data and scripts will be shared in public after publication.

## Contributions

Z.C.C. conceptualized the study. C.T.W implemented the experiment and collected the data.

Y.T.H performed data analysis and wrote the paper. Z.C.C and C.T.W edited the paper. All authors contributed to and approved the final paper.

## Supporting information

All supplementary materials

## Acknowledgments

We thank Junko Taniai and Mika Matsuo for helping with participant recruitment and experiment preparation, Dr. Iver Hsieh for website development and maintenance, and Dr. Felix B. Kern for acquiring the GPT scores. This study was supported by World Premier International Research Center Initiative (WPI), MEXT, Japan (to Z.C.C.), IRCN-Daikin SCP (to Z.C.C.), and Japan Society for the Promotion of Science, Japan (to Y.T.H).

