## Supplementary material for "Creativity Potential Networks: Brain Markers for Novelty and Feasibility of Upcoming Divergent Thinking Solutions": All supplementary materials

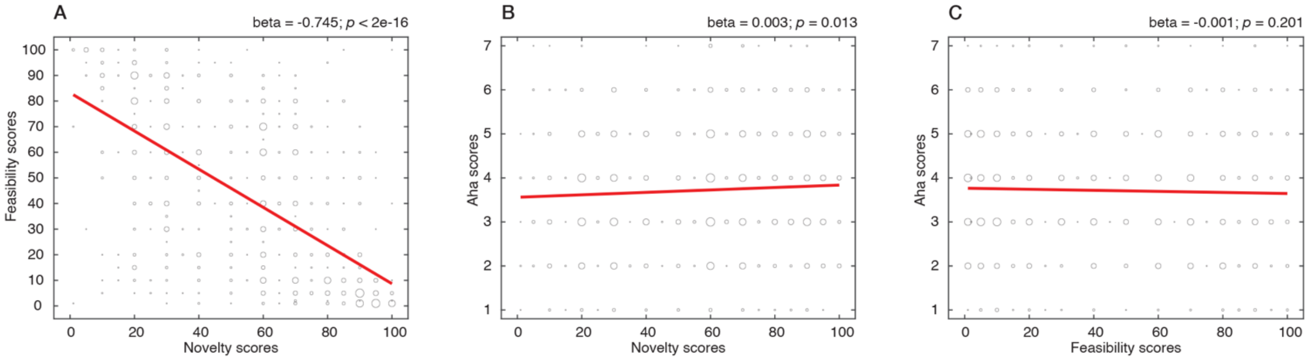


**Supplementary Figure 1. Visualization of correlations among novelty scores, feasibility scores, and Aha rating.** The size of the black hole circles increases with a higher number of occurrences of the score across trials and participants. The red line represents the predicted estimation derived from the linear mixed-effects model.

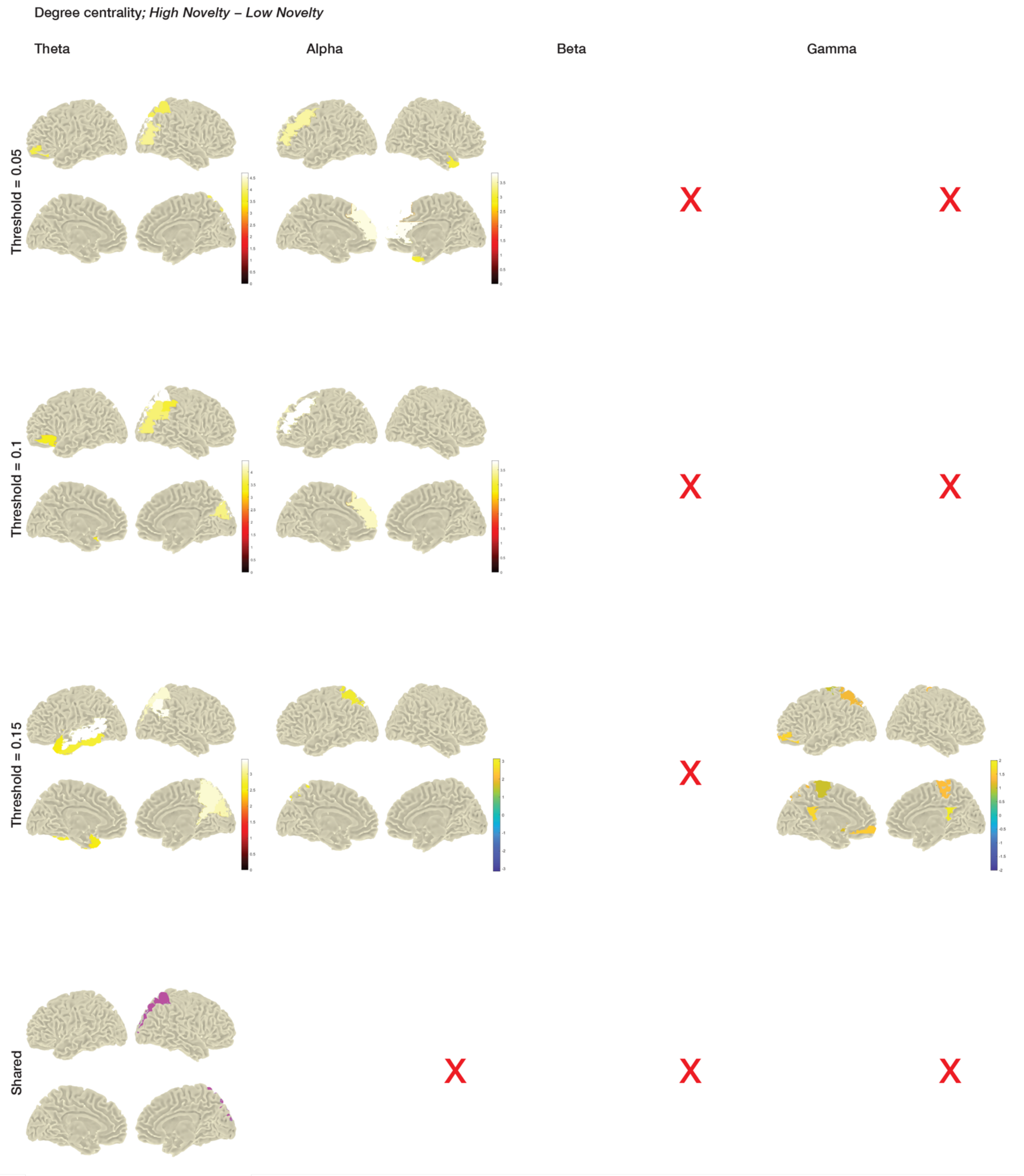


**Supplementary Figure 2.**
The color bars in the first, second and third rows represent statistical values.

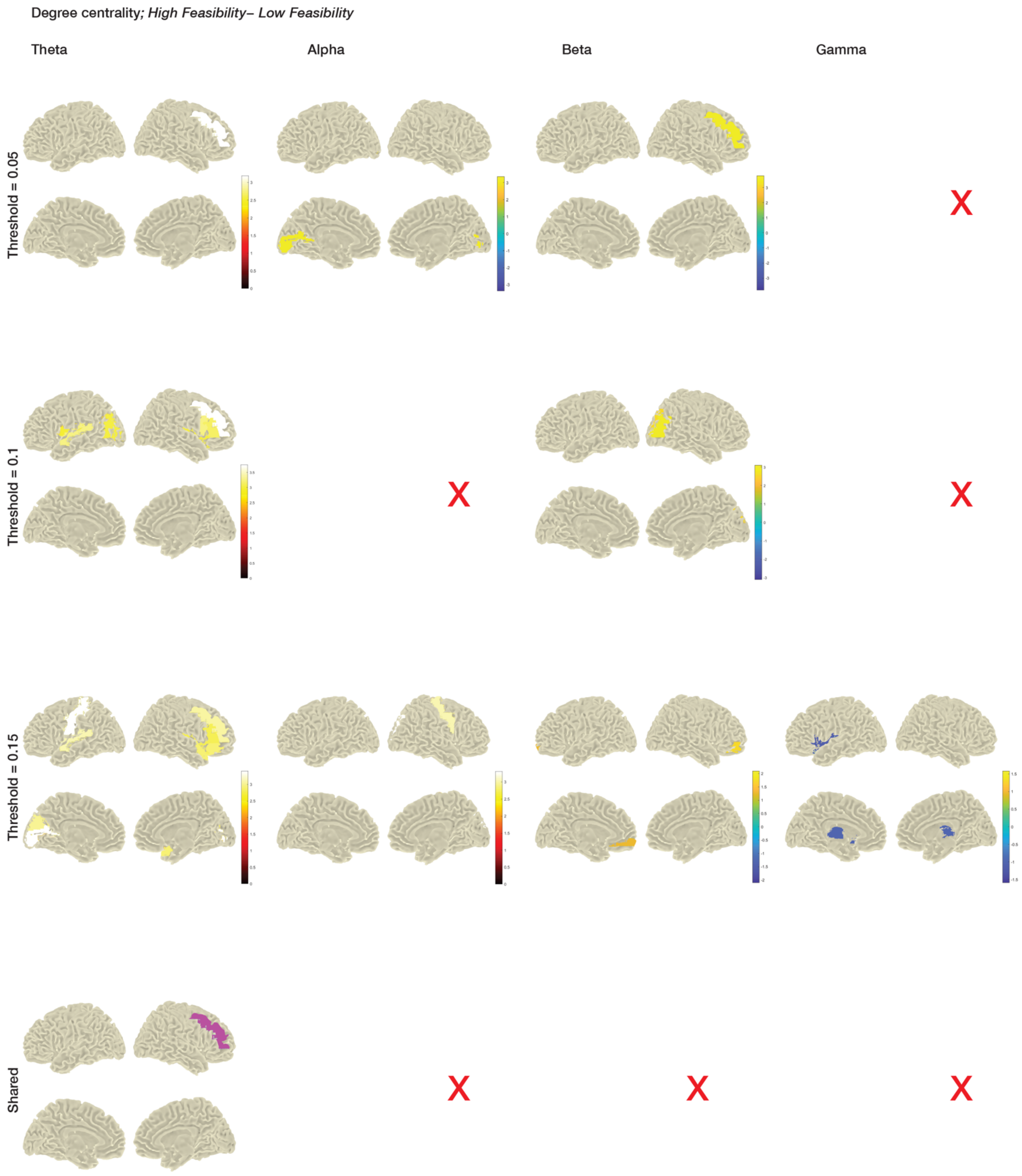


**Supplementary Figure 3.**

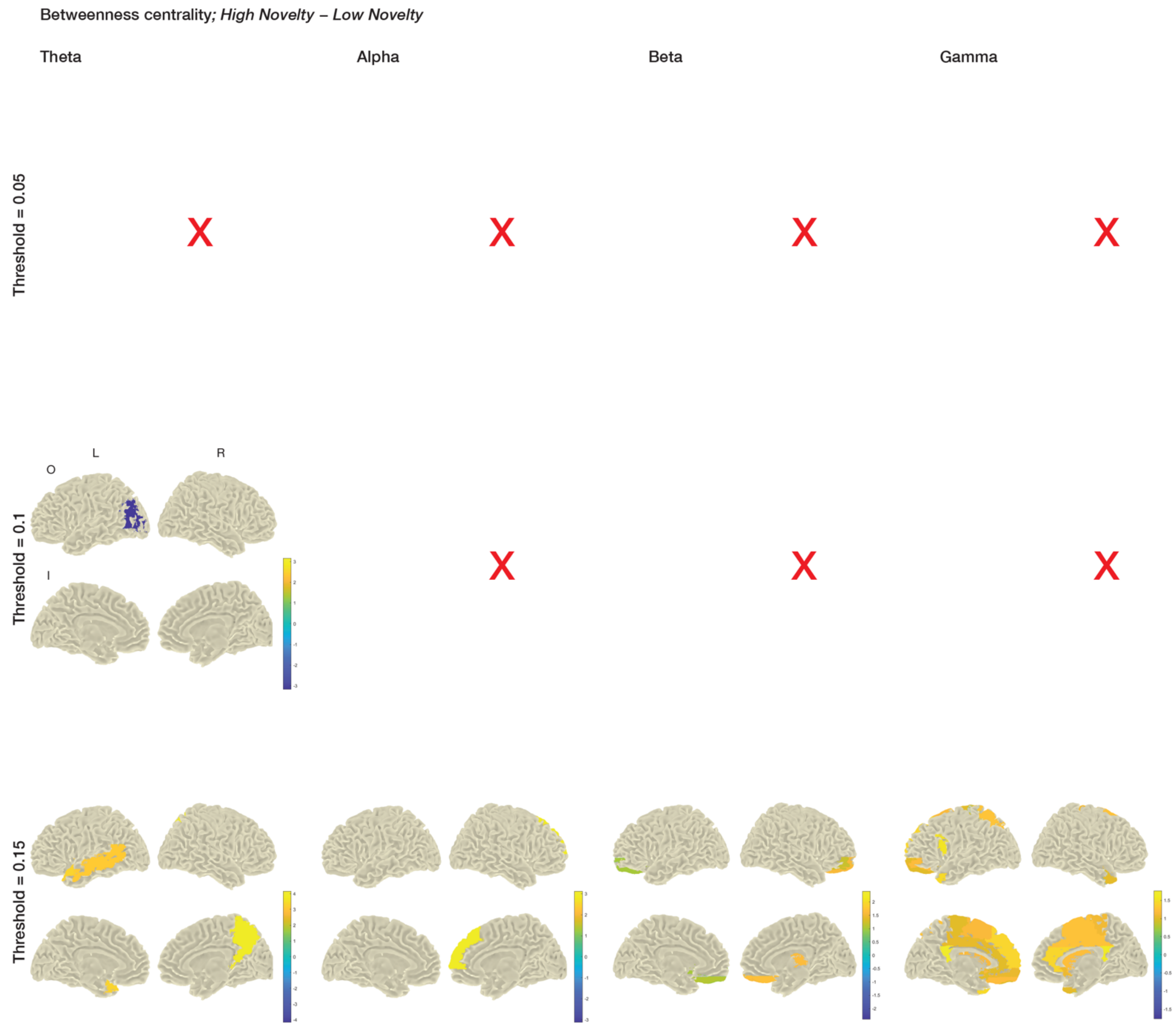


**Supplementary Figure 4.**

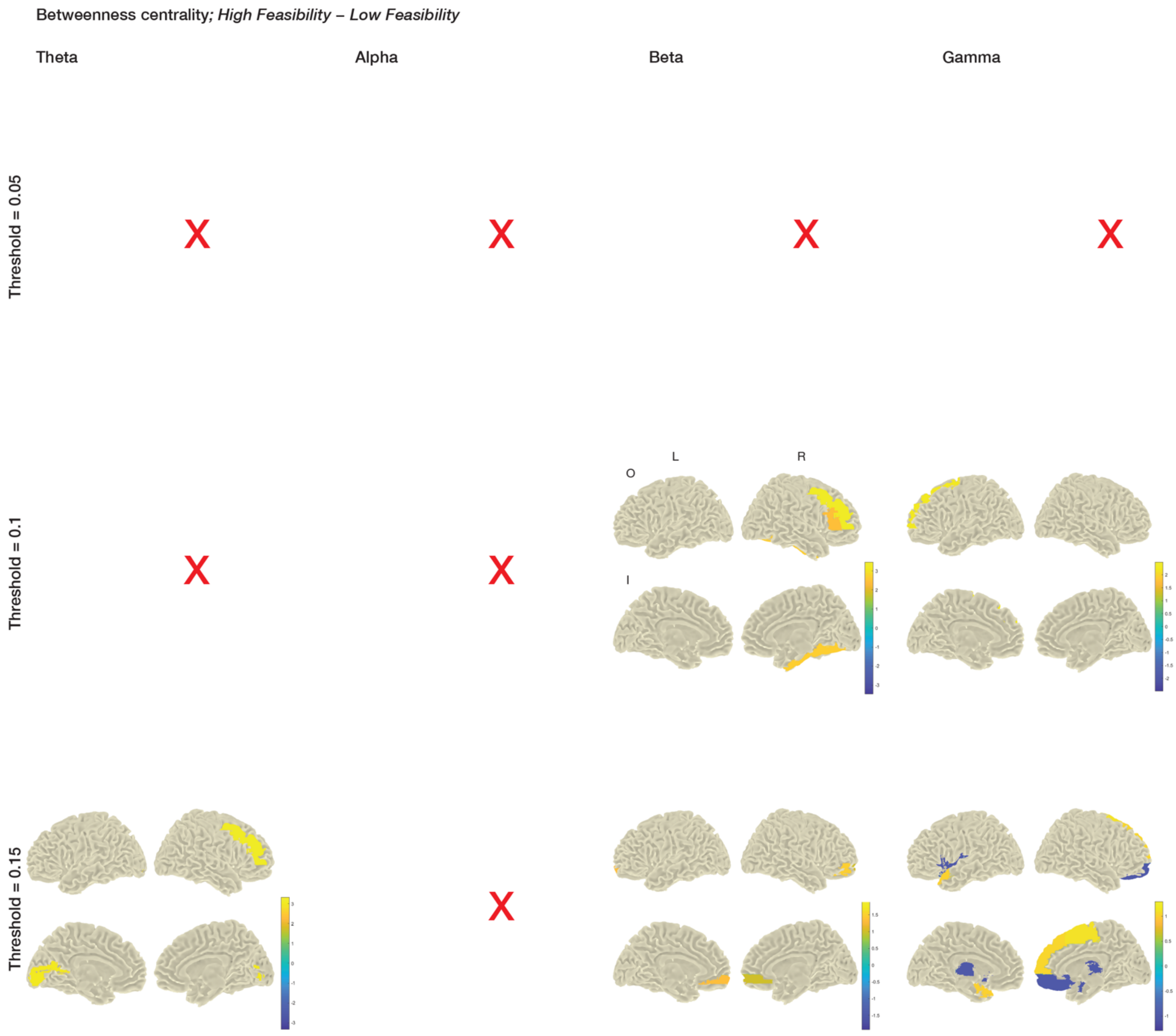


**Supplementary Figure 5.**

**Supplementary Table 1.**

| No. | Items in English |
|---|---|
| 1 | Brick |
| 2 | Umbrella |
| 3 | Ketchup |
| 4 | Rubber band |
| 5 | Newspaper |
| 6 | Paper cup |
| 7 | PET bottle |
| 8 | Slippers |
| 9 | Salt |
| 10 | Wood chopsticks |
| 11 | Drinking straw |
| 12 | Plastic bag |
| 13 | Handkerchief |
| 14 | Needle |
| 15 | T-shirt |
| 16 | Shoelace |
| 17 | Metal key |
| 18 | Candle |
| 19 | Toilet paper |
| 20 | Tape |
| 21 | Magnifier |
| 22 | Ball pen |
| 23 | Marble |
| 24 | CD |
| 25 | Carton |
| 26 | Cork |
| 27 | Credit card |
| 28 | Flashlight |
| 29 | Toothbrush |
| 30 | Coin |
